# *In vivo* antibody labeling route and fluorophore dictate labeling efficiency, sensitivity, and longevity

**DOI:** 10.1101/2024.08.10.607414

**Authors:** Natalie B. Hagan, Charles Inaku, Nikesh Kunder, Tayleur White, Thierry Iraguha, Anna Meyer, Kristen E. Pauken, Jason M. Schenkel

## Abstract

Leukocytes migrate through the blood and extravasate into organs to surveil the host for infection or cancer. Recently, we demonstrated that intravenous (IV) anti-CD45.2 antibody labeling allowed for precise tracking of leukocyte migration. However, the narrow labeling window can make this approach challenging for tracking rare migration events. Here, we show that altering antibody administration route and fluorophore can significantly extend the antibody active labeling time. We found that while both IV and intraperitoneal (IP) anti-CD45.2 antibody labeled circulating leukocytes after injection, they had different kinetic properties that impacted labeling time and intensity. Quantification of circulating antibody revealed that while unbound IV anti-CD45.2 antibody rapidly decreased, unbound IP anti-CD45.2 antibody increased over one hour. Using *in vitro* and *in vivo* serial dilution assays, we found that Alexa Fluor 647 (AF647) and Brilliant Blue 700 (BB700) dyes had the greatest labeling sensitivity compared to other fluorophores. However, IP antibody injection with anti-CD45.2 BB700, but not AF647, resulted in continuous blood leukocyte labeling for over 6 hours. Finally, we leveraged IP anti-CD45.2 BB700 antibody to track slower migrating leukocytes into tumors. We found that IP anti-CD45.2 antibody injection allowed for the identification of ∼seven times as many tumor-specific CD8^+^ T cells that had recently migrated from blood into tumors. Our results demonstrate how different injection routes and fluorophores affect anti-CD45.2 antibody leukocyte labeling and highlight the utility of this approach for defining leukocyte migration in the context of homeostasis and cancer.

## INTRODUCTION

To protect the host from homeostatic perturbations, including infection and cancer, leukocytes must migrate from the blood into tissue^1^. Leukocyte trafficking in both homeostasis and inflammation is a highly orchestrated process, involving combinations of selectins, chemokine receptors, and integrins^2–4^. After binding their requisite ligands, immune cells roll and arrest on vascular endothelium before extravasating and entering tissue^1^. While these migration mechanisms are well defined in multiple contexts, identifying leukocytes that had recently trafficked to tissue has, up until recently^5–8^, remained a significant challenge in the field.

Prior work examining the rate at which immune cells migrate into tissues have used a variety of different techniques and modalities. The gold standard approach for examining leukocyte turnover in tissues has been parabiosis, which involves making mirrored skin incisions on the flanks of two mice, and then physically conjoining animals using sutures or surgical staples^9^. Over the course of several days, the vasculature fuses between the two mice and blood leukocytes equilibrate over the course of roughly one week^9^. Alternatively, photoconvertible protein reporters, which change fluorescence properties for a defined period of time after laser excitation, have also allowed for the assessment of leukocyte replacement and egress rates in laser accessible tissue locations^10–12^. However, both methodologies have significant drawbacks. Recent work over the last several years has demonstrated that intravenous (IV) injection of anti-CD45 antibodies in non-human primates and mice allows for tracking of leukocyte migration from blood to tissues^5–8^. Indeed, we recently demonstrated that in mice, IV anti-CD45.2 antibody 1) actively bound to cells *in vivo* for roughly 45 minutes, 2) remained cell associated for approximately 72 hours, 3) did not leak from the vasculature into tissue, 4) could be injected multiple (>5) times without blocking or saturating available CD45.2, and 5) did not deplete CD45.2^+^ cells or change their activation status^7^. We previously used this technique to identify migration of leukocytes into tumors. However, while this method allowed for tracking leukocyte populations that rapidly entered tissues, rarer migration events were difficult to capture due to a limited active binding window. Therefore, we interrogated whether modifications could be made to delivery route and/or fluorophore to further optimize the labeling technique and expand the labeling window, enabling the detection of greater numbers of trafficking leukocytes.

The peritoneal cavity is an important anatomic site, containing numerous visceral organs, immune cells, and serous fluid that helps to maintain organ homeostasis^13^. Research over many decades has demonstrated that protein or pathogen injection into the peritoneal cavity results in systemic dissemination through the vasculature^14,15^. This is not due to direct absorption from the peritoneal space into the blood, but rather due to lymphatic drainage into the thoracic duct^14,16^. Indeed, using fluorescent tracers, previous work demonstrated that compounds injected into the peritoneum were found in both paravertebral and diaphragmatic lymphatic vessels^16^. More recently it was shown that initial drainage was mediated by specialized capillary mesenteric lymphatic vessels that bypassed mesenteric lymph nodes and shuttled peritoneal fluid directly into the mediastinal lymph node before emptying into the thoracic duct^17^. Because compounds injected into the peritoneal cavity must drain through lymphatics and the mediastinal lymph node, there is a temporal delay before they can be found in blood^14^. Concordantly, antibody injected into the peritoneal cavity slowly increases in concentration over time in mice^14^. These data suggest that antibody injection into the peritoneal cavity may represent an alternative route to slowly release anti-CD45.2 antibody into blood over time, potentially increasing the window to bind circulating leukocytes.

Here, we interrogated how to increase the *in vivo* antibody labeling window to track low frequency migration events. First, we examined whether route of antibody injection altered labeling kinetics. We found that IV anti-CD45.2 antibody injection rapidly (within one minute) labeled blood contiguous T cells in the spleen, bone marrow, and liver. On the other hand, intraperitoneal (IP) injection of anti-CD45.2 antibody did not label peripheral blood immune cells until 10 minutes after injection. Moreover, the percent of anti-CD45.2 antibody positive cells and the geometric mean fluorescence intensity (gMFI) on IP CD45.2 antibody labeled cells generally increased over the course of one hour, suggesting continuous labeling during this time. We then quantified the amount of free, circulating anti-CD45.2 antibody after IV versus IP injection and found that IV anti-CD45.2 antibody levels rapidly plummeted after injection, becoming almost undetectable after one hour. However, IP anti-CD45.2 antibody injection gradually increased over time, reaching maximal plasma concentration one-hour post-injection, suggesting that by injecting antibody IP, there was a slow release of antibody into the blood. We next determined which fluorophores allowed for the greatest level of labeling at the lowest concentrations by performing serial dilutions with anti-CD45.2 antibody *in vitro* or *in vivo* and found that Alexa Fluor 647 (AF647) and Brilliant Blue 700 (BB700) were the most robust in terms of labeling immune cells. Further *in vivo* testing revealed that anti-CD45.2 BB700, unlike anti-CD45.2 AF647, continued to robustly label circulating immune cells six hours after IP antibody injection. Finally, we found that IP anti-CD45.2 BB700 antibody labeled roughly 7-fold more tumor-infiltrating, antigen specific CD8^+^ T cells than IV anti-CD45.2 AF488 antibody 24 hours after injection. Collectively these data highlight the importance of antibody injection route and fluorophore selection, which, when optimized, can reveal rare migration events *in vivo*.

## MATERIALS AND METHODS

### Mice and Tumor Injection

Naïve C57BL/6 mice aged 7-16 weeks were used for all studies. Experimental mice were housed under SPF conditions. For B16-OVA tumor studies, mice were anesthetized with isoflurane. Flank skin was shaved with clippers, cleaned with 70% ethanol, and then 2.5×10^5^ B16-OVA melanoma cells were injected subcutaneously. All studies were performed under an animal protocol approved by the MD Anderson Cancer Center IACUC. Mice were assessed for morbidity and pain according to MD Anderson Cancer Center IACUC guidelines and humanely sacrificed prior to natural expiration.

### Cell Lines

B16-OVA were purchased from Millipore. B16-OVA were cultured in DMEM without pyruvate and supplemented with 1%penicillin/streptomycin and 10% fetal bovine serum (FBS).

### Plasma Collection for ELISA

Naïve C57BL/6 mice between 7-10 weeks old were injected IV with anti-CD45.2 antibody conjugated to PE and IP anti-CD45.2 antibody conjugated to APC. Mice were then bled with a glass Pasteur pipette containing 4% sodium citrate at either 1-, 10-, 30-, or 60-minutes post antibody injection. Blood samples were placed in microcentrifuge tubes and spun down at 2000 RCF at 4°C for 10 minutes to separate plasma from cells. Plasma supernatant was collected and diluted in PBS at a 1:8 diluton.

### Anti-CD45.2 antibody ELISA

96-well flat bottom plates were coated with 100 μL of purified anti-phycoerythrin or purified anti-allophycocyanin at a concentration of 6 μg/mL overnight at 4°C. The next day, plates were washed three time with 300 μL of wash buffer (0.05% Tween20 in 1X PBS) and were subsequently blocked with 100 μL of 10% BSA in 1X PBS for one hour covered at room temperature. After one hour, plates were washed with wash buffer three times. Afterwards, 100 μL of either plasma or standard was added to each well. The standard and plasma samples were incubated for one hour at room temperature. After one hour, plates were washed with wash buffer three times, and then 100 μL of recombinant rabbit anti-mouse IgG2c antibody conjugated to biotin at a concentration of 12.5 ng/mL in 1X PBS was added to each well. After one hour, plates were washed with wash buffer three times. Next, 100 μL of HRP-Streptavidin at a concentration of 100 ng/mL was added to each well and the plate was incubated at room temperature for one hour. After one hour, plates were washed with wash buffer three times, and then 100 μL of TMB solution (1X) was added to each well for ∼20-30 minutes at room temperature or until desired blue color was achieved. The reaction was stopped with 100 μL of stop solution was added to end the enzymatic reaction. Absorbances were then read at 450nm in a standard plate reader.

### Isolation/Preparation of Cell Suspensions

Spleens and livers were dissociated by using syringe backs on 70-micron filters over 50 mL conical tubes. The 70-micron filters were rinsed with RPMI 1640 supplemented with 5% heat-inactivated FBS. Cell suspensions were spun down at 2,000 RPM for 5 minutes. Cells were then resuspended in ACK lysis buffer to eliminate red blood cells (RBCs) and incubated for one minute before being quenched with 5 mL of PBS supplemented with 1% heat-inactivated fetal bovine serum. Peripheral blood was collected by retro-orbital bleeds into tubes with 100 μL of 4% sodium citrate. Blood was then spun down at 2,000 RPM for 5 minutes and resuspended in 1 mL of ACK lysis buffer to eliminate RBCs. Blood was incubated for 1 minute before quenching with 5 mL of PBS supplemented with 1% FBS. This was repeated a second time to remove excess RBCs. Bone marrow was collected by taking one femur, cutting off one end, and spinning the bone cut side down into a 1.5 mL Eppendorf with 300 ml of RPMI 1640 supplemented with 5% FBS. Bone marrow was then spun down at 2,000 RPM for 5 minutes and resuspended in 1 mL of ACK lysis buffer to eliminate RBCs. Blood was incubated for 1 minute before quenching with 5 mL of PBS supplemented with 1% FBS.

### Staining for Flow Cytometric Analysis

Approximately 1-3×10^6^ cells were stained for 30 minutes at 4°C in 96-well V-bottom plates in FACS buffer (DPBS, 1x without calcium and magnesium supplemented with 1% heat inactivated FBS). After staining, cells were washed two times with 200 μL of FACS buffer and then subsequently fixed with a Fixation/Permeabilization kit (eBioscience). Tumor-specific CD8^+^ T cells were assessed using p15e_604-611_(KSPWFTTL)/H2-K^b^ tetramer, which was prepared using streptavidin-PE (Invitrogen) and p15e/H-2K^b^ monomer from the NIH Tetramer Core. The following antibody clones from the indicated companies were used for this study: Biolegend – CD11b (M1/70), Ly6G (1A8), Ly-6C (HK1.4), B220 (RA3-6B2), CD45.2 (104), CD4 (RM4-5), CD8a (53-6.7), PD-1 (RMP1-30), CD44 (IM7). For CFSE and Tag-IT Violet, staining was performed as indicated by Biolegend.

### *In vivo* labeling

For longitudinal intravenous labeling of circulating leukocytes, mice were injected with anti-CD45.2 antibody conjugated to Alexa Fluor 488, Alexa Fluor 594, Alexa Fluor 647, Brilliant Violet 510, Brilliant Violet 605, and Brilliant Blue 700. A total of 5 μg of each antibody was injected IV or IP as indicated in a volume of 100 μL.

## RESULTS

### IP antibody labeling results in a delayed and gradual in vivo labeling compared to IV antibody

To determine circulating leukocyte labeling kinetics *in vivo*, naïve C57BL/6 mice were co-injected IV with 5 μg of anti-CD45.2 phycoerythrin (PE) antibody and IP with 5 μg of anti-CD45.2 allophycocyanin (APC) antibody (Fig. 1A). One minute after injection, all circulating leukocytes, including CD4^+^ T cells, were IV anti-CD45.2 PE^+^ (Fig. 1B, Fig. S1A-E). Virtually no leukocytes were labeled with IP anti-CD45.2 APC antibody. However, 10 minutes after IV and IP anti-CD45.2 antibody injection, greater than 99% of circulating CD4^+^ T cells were labeled with both IP and IV anti-CD45.2 antibody (Fig. 1B, Fig. S1A-E). APC is a protein-based fluorophore that has a molecular weight of approximately 110 kilodaltons (kDa), or about two thirds the size of an average IgG antibody. Thus, it was possible that the delayed IP labeling kinetics was due to APC increasing the molecular weight of anti-CD45.2 antibody, necessitating extra time to drain from the peritoneal cavity. To test this hypothesis, we co-injected 5 μg of anti-CD45.2 APC IP with 5 μg of anti-CD45.2 Alexa Fluor 488 (AF488). AF488 is ∼0.6 kDa, or less than 1/200^th^ the molecular weight of APC^18^. We found that IP anti-CD45.2 AF488 antibody labeling was virtually identical to IP anti-CD45.2 APC, demonstrating that APC fluorophore molecular weight was not the reason for the delayed labeling seen with IP antibody injection (Fig. S1F).

**Fig 1.**
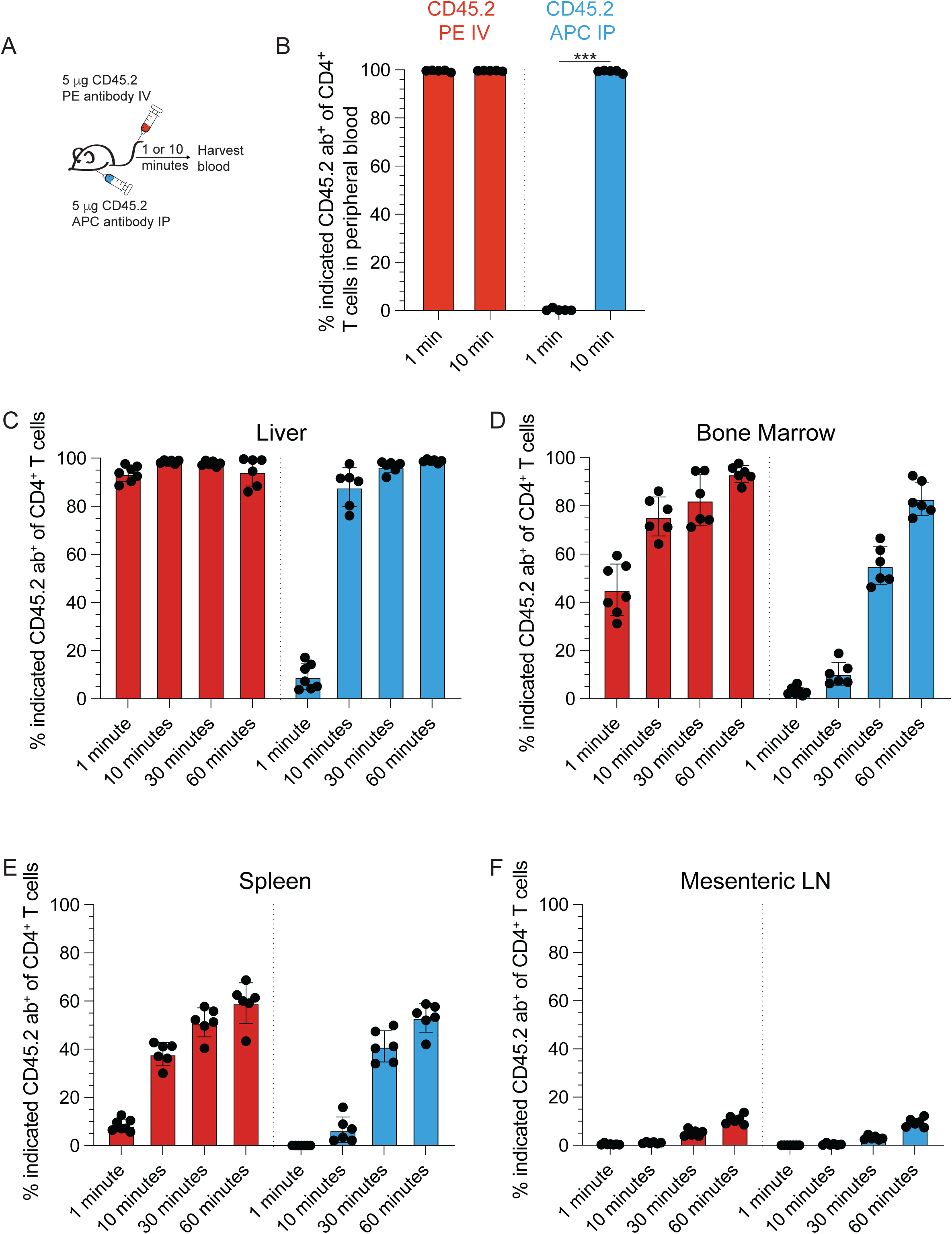
IP anti-CD45.2 antibody labeling is delayed compared to IV anti-CD45.2 antibody labeling and occurs over at least one hour. (A&B) C57BL/6 mice were co-injected with 5 μg of IV anti-CD45.2 PE and 5 μg of IP anti-CD45.2 APC. Mice were bled one or ten minutes after IP and IV injection. (A) Experimental schematic. (B) Percent of peripheral blood CD4+ T cells positive for IV anti-CD45.2 PE or IP anti-CD45.2 APC either one or ten minutes after injection. (C-F) C57BL/6 mice were co-injected with 5 μg of IV anti-CD45.2 PE and 5 μg of IP anti-CD45.2 APC. Mice were then euthanized either 1, 10, 30, or 60 minutes after injection, and liver, bone marrow, spleen and mesenteric lymph nodes were harvested and processed for flow cytometry. Percent of CD4^+^ T cells positive for IV anti-CD45.2 PE or IP anti-CD45.2 APC after 1, 10, 30 or 60 minutes in (C) Liver, (D) Bone Marrow, (E) Spleen or (F) Mesenteric lymph node; blue bars = IP anti-CD45.2 APC and red bars = IV anti-CD45.2 PE. Each dot represents a mouse. Data shown are representative of 2-3 independent experiments, with a total of 6-8 mice per group. Error bars represent SD. Student’s t-test was used for (B) and One way ANOVA was performed to determine statistical significance (C-F). Statistical significance for (C-F) can be found in supplemental table 1. ***=p<0.001.

For many organs, the vascular endothelium represents an important and efficient barrier that allows for gas and nutrient exchange without direct leak of circulating cells or plasma in homeostasis^19^. However, there are multiple organs, including splenic red pulp, liver, and bone marrow, where the endothelium contains gaps/fenestra or is discontinuous, allowing blood to infiltrate directly into the tissue^19^. Hence, for the purposes of *in vivo* antibody labeling, these organs represent blood contiguous compartments, meaning that immune cells found in these tissues should rapidly label with anti-CD45.2 antibody after injection. Thus, we next examined the percentage and fluorescence intensity of labeled immune cells after IV anti-CD45.2 PE and IP anti-CD45.2 APC at 1-, 10-, 30-, or 60-minutes post-injection in blood contiguous compartments. We used the mesenteric lymph node as a control, as antibody does not freely leak into lymph node after injection^7^. Here, we found that IV anti-CD45.2 PE rapidly labeled liver, bone marrow, and spleen, with observable differences in both rate and fluorescence intensity (Fig. 1C-F, S1G). Specifically, liver CD4^+^ T cells labeled the most rapidly, while CD4^+^ T cells in both the bone marrow and spleen labeled slower and with a much lower fluorescence intensity for IV anti-CD45.2 antibody. This decrease in labeling efficiency is probably due to local leukocyte density being higher in both splenic red pulp and bone marrow compared to liver, as well as the fact that the liver is likely exposed to IV anti-CD45.2 antibody almost immediately after IV injection. On the other hand, IP anti-CD45.2 APC antibody labeling of splenic red pulp, liver, and bone marrow was, as expected, delayed compared to IV anti-CD45.2 PE antibody (Fig. 1C-F, S1H). Indeed, much like peripheral blood, CD4^+^ T cells in the liver did not demonstrably label until 10 minutes after IP injection. Label intensity increased from 10 to 30 to 60 minutes. Moreover, a greater temporal delay was noted in the spleen and bone marrow, and these CD4^+^ T cells labeled with less IP anti-CD45.2 APC antibody compared to liver CD4^+^ T cells. Taken together, these data demonstrate the rapid nature of IV anti-CD45.2 antibody labeling and reveal the more delayed labeling seen with IP anti-CD45.2 antibody injection.

### Route of injection dictates kinetics and concentration of unbound antibody in plasma

Because we saw a continuous increase in IP anti-CD45.2 APC fluorescence intensity over the course of an hour after injection, we hypothesized that antibody was gradually draining from the peritoneal cavity. Thus, we performed an ELISA to quantify unbound IV and IP anti-CD45.2 antibody in the plasma to determine the concentration of unbound antibody over time. C57BL/6 mice were co-injected with 5 μg of IV anti-CD45.2 PE and 5 μg of IP anti-CD45.2 APC, and mice were either bled 1-, 10-, 30-, or 60-minutes after injection. Quantification of IV anti-CD45.2 PE revealed high concentrations of unbound antibody at 1-minute post-injection (Fig. 2A). Ten minutes after injection, there was a ∼35% decrease in plasma concentration, and by 30- and 60-minutes post-injection the concentration had decreased by ∼6-fold and ∼12-fold respectively (Fig. 2A). On the other hand, IP anti-CD45.2 APC was undetectable at 1-minute post-injection, and gradually increased over the course of one hour (Fig. 2A). Moreover, the concentration of IP anti-CD45.2 APC remained relatively low compared to the initial IV anti-CD45.2 PE antibody (Fig 2A). To determine if the concentration remained low because of immediate binding to circulating leukocytes, we co-injected CD45.1^+^ C57BL/6 mice with both IV anti-CD45.2 PE and IP anti-CD45.2 APC antibodies. While IV anti-CD45.2 PE antibody initially dropped and stabilized at ∼1 ug/mL in plasma, IP anti-CD45.2 APC still gradually increased over time, demonstrating the slow release of antibody from the peritoneal cavity (Fig. 2B). Taken together, these data reveal the gradual accumulation of accumulation of unbound IP anti-CD45.2 antibody in plasma.

**Fig 2.**
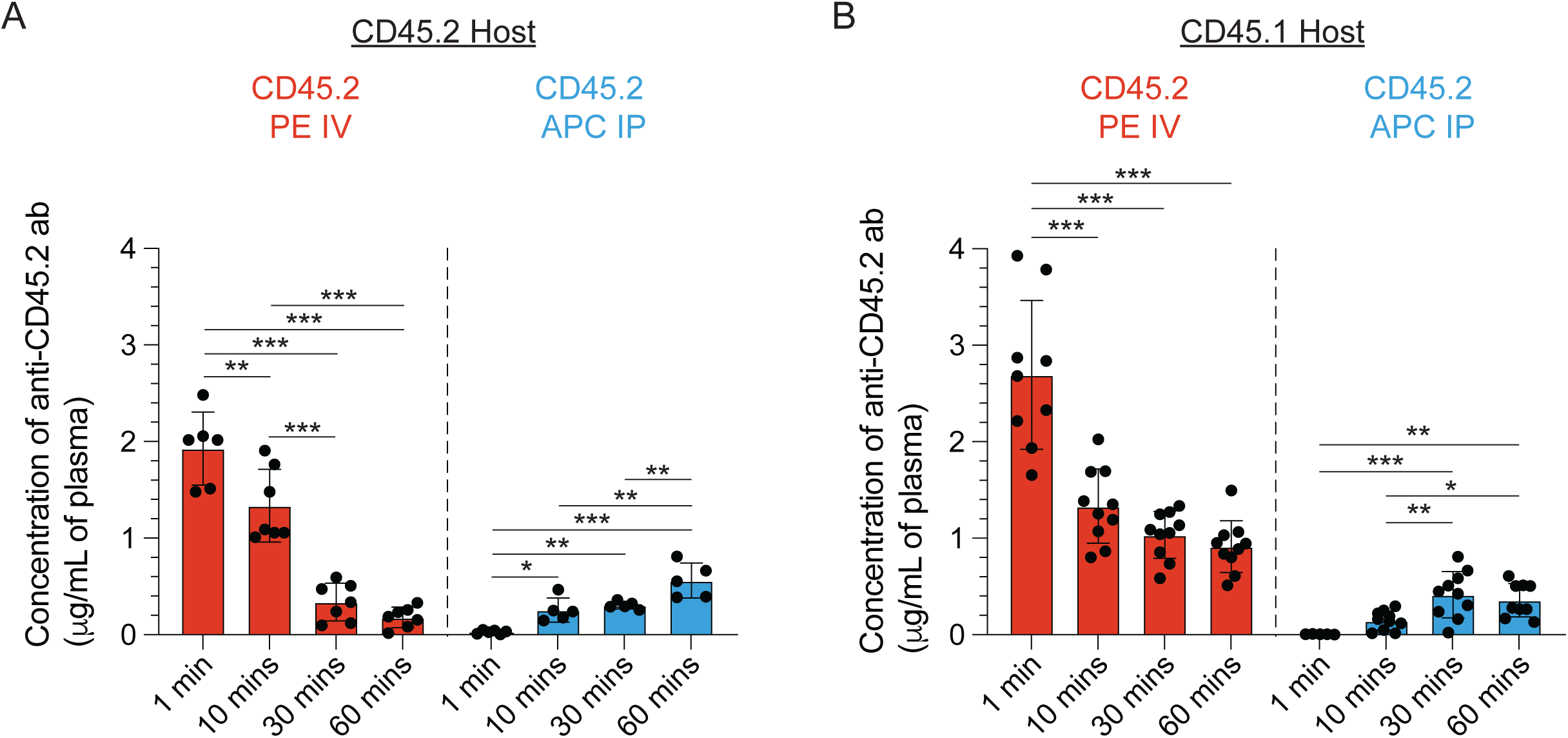
Unbound, plasma IV anti-CD45.2 antibody concentration rapidly decrease, while IP anti-CD45.2 antibody concentration steadily increases over time. (A) C57BL/6 or (B) CD45.1^+^ C57BL/6 mice were co-injected with IV anti-CD45.2 PE or IP anti-CD45.2 APC and were bled 1-,10-,30-, or 60-minutes post injection. Plasma was separated from blood by centrifugation and IV anti-CD45.2 PE and IP anti-CD45.2 APC were quantified by ELISA. Each dot represents a mouse. Data shown are representative of 2-3 independent experiments, with a total of 6-8 mice per group. Error bars represent SD. One way ANOVA was performed to determine statistical significance. *=p<0.05, **=p<0.01, ***=p<0.001.

### Fluorophore selection is a critical determinant of labeling sensitivity in vitro and in vivo

Because IP anti-CD45.2 antibody continued to circulate as free antibody at low concentrations for over an hour, we next determined which fluorophores were able to label leukocytes at small antibody concentrations. We focused on anti-CD45.2 antibodies conjugated to fluorophores we have previously used *in vivo*, including AF488, Alexa Fluor 594 (AF594), Alexa Fluor 647 (AF647), Brilliant Violet 510 (BV510), Brilliant Violet 605 (BV605)^7^, and also added a new conjugate, Brilliant Blue 700 (BB700). We first tested the ability of these anti-CD45.2 antibodies conjugated to different fluorophores to label naive splenocytes *in vitro* at different concentrations. We found that all fluorophores performed well *in vitro* even at total amounts as low 1.5625 ng of anti-CD45.2 antibody (Fig. 3A).

**Fig 3.**
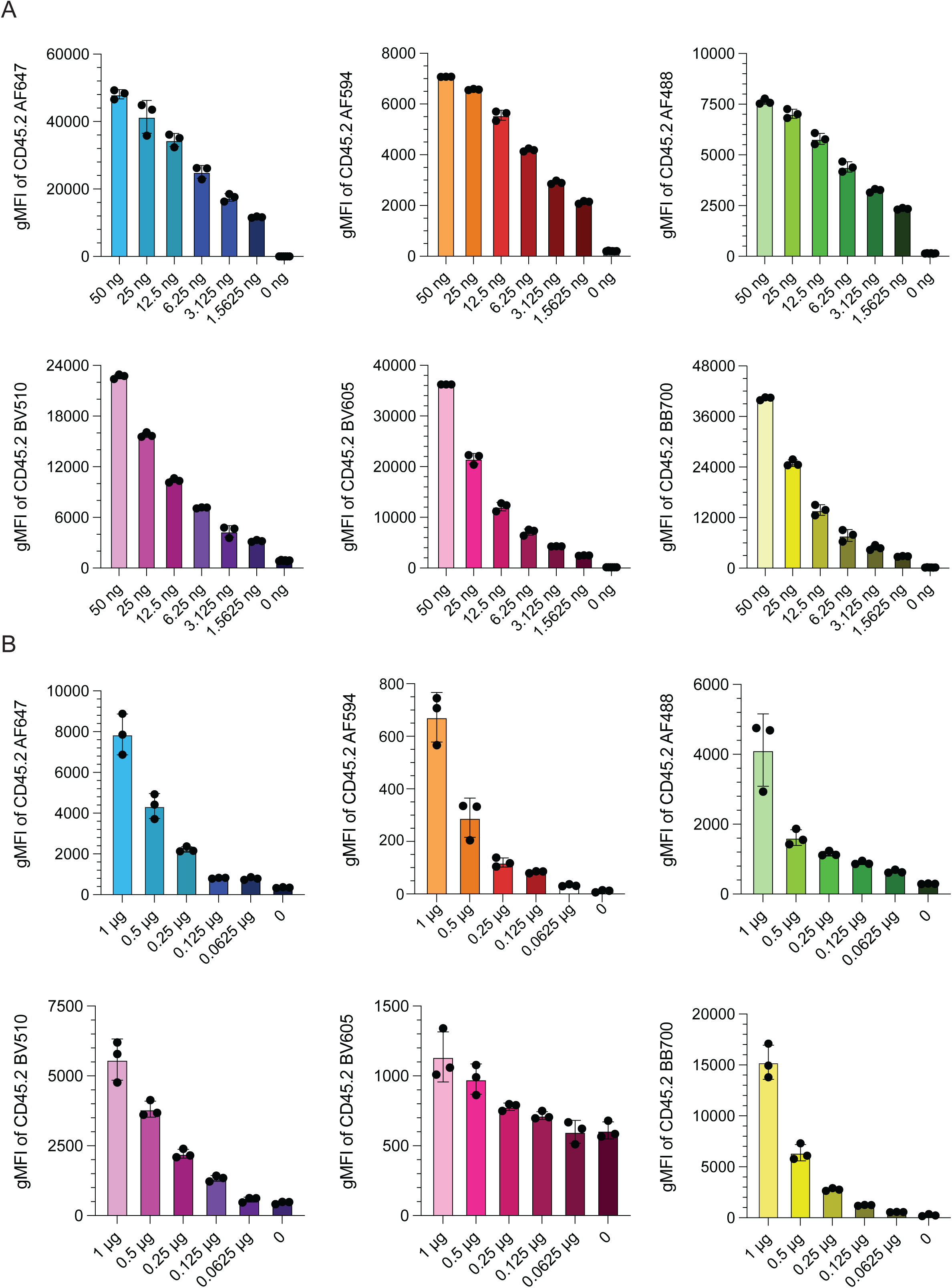
*In vitro* and *in vivo* anti-CD45.2 antibody conjugated to different fluorophore titrations. (A) 1×10^6^ splenocytes were incubated with the indicated amount of anti-CD45.2 antibody conjugated to the indicated fluorophores for 1 hour at 37 °C degrees and fluorescence intensity was evaluated by flow cytometry. (B) 20×10^6^ Tag-IT labeled splenocytes were injected IV and the indicated amount of anti-CD45.2 antibody with the indicated fluorophore were immediately injected IV. One hour after injection, mice were euthanized, spleens were harvested and processed for flow cytometry. Data shown is fluorescence intensity of the indicated fluorophore for anti-CD45.2 antibody on transferred Tag-IT^+^ splenocytes. Each dot represents a mouse. Data shown are representative of 2-3 independent experiments, with a total of 6-8 mice per group. Error bars represent SD. One way ANOVA was performed to determine statistical significance. Statistical significance can be found in supplemental table 1. ***=p<0.001.

Next, we determined the lowest amount of anti-CD45.2 antibody fluorophore that could be injected to label circulating leukocytes *in vivo*. Our previous work has demonstrated that as little as 1 μg of anti-CD45.2 antibody is sufficient to fully label circulating leukocytes in peripheral blood. Thus, we started our titration with 1 μg as the positive control and decreased the amount of injected antibody by two-fold, with the lowest dose being 0.0625 μg of anti-CD45.2 antibody. To control for labeling efficiency due to alterations in antibody concentration, we batched all antibodies together (e.g. 1 μg of fluorophore A, 0.5 μg of fluorophore B, 0.25 μg of fluorophore C, 0.125 μg of fluorophore D, and 0.0625 μg of fluorophore E). Tag-IT Violet labeled splenocytes were injected into naïve C57BL/6 mice and then anti-CD45.2 antibody fluorophore mixtures were immediately injected IV. One hour after antibody injection, mice were euthanized, and fluorescence intensity was examined on transferred Tag-IT violet splenocytes. While all fluorophores performed well at 1 μg doses, anti-CD45.2 antibody conjugated to BV605 rapidly lost sensitivity at a dose of 0.5 μg (Fig. 3B). This was followed up by anti-CD45.2 antibody conjugated to AF594 and AF488 (Fig. 3B). Anti-CD45.2 antibody conjugated to BV510, AF647, and BB700 performed the best at the lowest doses, however BV510 also had substantially higher fluorescence background compared to AF647 and BB700, making it less suitable for low dose *in vivo* labeling (Fig. 3B).

### Anti-CD45.2 antibody conjugated to BB700 continuously labels circulating cells for several hours and lasts for up to three days

Since our previous experiments identified AF647 and BB700 as the most sensitive to label circulating immune cells at low concentrations, we next determined how long each of these anti-CD45.2 fluorophore antibodies would label leukocytes *in vivo* after IP injection. C57BL/6 mice were injected with CFSE-labeled splenocytes IV, and then were immediately injected IP with either anti-CD45.2 antibody conjugated to AF647 or BB700 (Fig. 4A). Six hours after antibody injection, Tag-IT Violet-labeled splenocytes were injected IV. Finally, 24 hours after antibody injection, mice were injected with dual CFSE and Tag-IT Violet labeled splenocytes. Two hours after injection, mice were euthanized and anti-CD45.2 BB700 and AF647 geometric mean fluorescence intensity and positivity was assessed on transferred splenocytes. Here, we found that both anti-CD45.2 AF647 and BB700 robustly labeled splenocytes transferred at the time of IP antibody injection (Fig. 4B-E). However, CD45.2 AF647 was greatly diminished on splenocytes transferred six hours after IP antibody injection in terms of both gMFI, as well as the percent of transferred splenocytes that were positive for anti-CD45.2 AF647 antibody (Fig. 4B&C). On the other hand, anti-CD45.2 BB700 antibody still robustly labeled splenocytes transferred six hours after IP injection (Fig. 4D&E). While BB700 gMFI decreased by roughly 50%, anti-CD45.2 BB700 antibody labeling remained on over 90% of splenocytes transferred 6 hours after IP antibody (Fig. 4D&E). Notably, neither anti-CD45.2 AF647 or BB700 labels could be identified on splenocytes transferred 24 hours after IP antibody injection (Fig. 4B-E).

**Fig 4.**
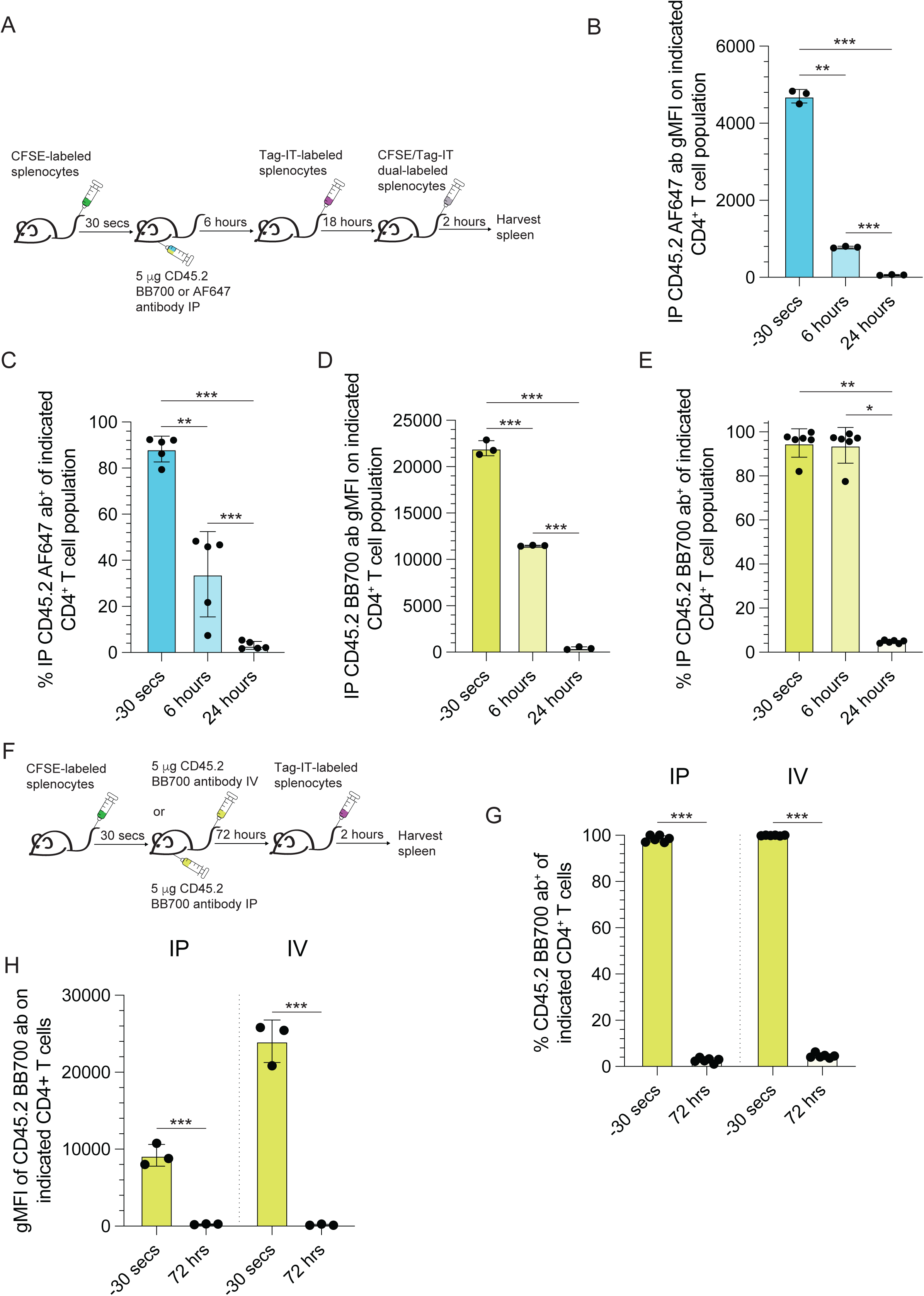
IP anti-CD45.2 BB700 continuously labels for six hours and remains detectable for up to three days. (A-E) C57BL/6 mice were IV injected with 20×10^6^ CFSE labeled splenocytes and then immediately injected IP with either 5 μg of anti-CD45.2 AF647 or BB700. Six hours later, mice were IV injected with 20×10^6^ Tag-IT violet labeled splenocytes. One day after antibody injection, mice were IV injected with 20×10^6^ CFSE/Tag-IT double labeled splenocytes and were euthanized two hours later and evaluated by flow cytometry (A) Experimental schematic for B-E. (B) Fluorescence intensity and (C) Percent positive of anti-CD45.2 AF647 on the indicated CD4^+^ T cell transferred population. (D) Fluorescence intensity and (E) Percent positive of anti-CD45.2 BB700 on the indicated CD4^+^ T cell transferred population. (F-H) C57BL/6 mice were IV injected with 20×10^6^ CFSE labeled splenocytes and then immediately injected with either 5 μg of IP or IV anti-CD45.2 BB700. Three days later, mice were IV injected with 20×10^6^ Tag-IT violet labeled splenocytes and were euthanized two hours later and evaluated by flow cytometry. (F) Experimental schematic for G and H. (G) Percent positive and (H) Fluorescence intensity of IP or IV anti-CD45.2 BB700 on transferred CD4^+^ T cells. Each dot represents a mouse. Data shown are representative of 2-3 independent experiments, with a total of 6-8 mice per group. Error bars represent SD. One way ANOVA was performed to determine statistical significance. *=p<0.05, **=p<0.01, ***=p<0.001.

Next, we assessed the longevity of IP anti-CD45.2 BB700 antibody labeling. Our previous work demonstrated that anti-CD45.2 antibody could be detected bound to leukocytes for up to three days^7^. Thus, we next compared how IP and IV anti-CD45.2 BB700 antibody compared to one another. C57BL/6 mice were injected IV with CFSE labeled splenocytes, and then thirty seconds later mice were injected with either IP or IV with anti-CD45.2 BB700 (Fig. 4F). Three days after antibody injection, Tag-IT Violet labeled splenocytes were injected IV as a negative control and then mice were euthanized two hours later. We found that transferred CFSE labeled splenocytes were uniformly positive for anti-CD45.2 BB700 antibody regardless of whether it was injected IP or IV (Fig. 4G). However, the BB700 gMFI was significantly lower on CFSE labeled splenocytes from IP injected mice compared to IV injected mice, suggesting that while IP was still able to label circulating immune cells well after three days, it did not drive as efficient of a labeling event as IV anti-CD45.2 BB700 (Fig. 4H).

### IP anti-CD45.2 BB700 antibody administration allows for improved identification of recently recruited CD8^+^ T cells to tumors

Our previous work demonstrated that multiple innate and adaptive immune lineages were rapidly recruited into several mouse tumor models, including B16 melanoma^7^. However, unlike many other leukocytes, tumor-specific CD8^+^ T cells migrated into tumors at a much slower rate. Thus, we next determined if IP anti-CD45.2 BB700 would enhance the ability to identify recently recruited tumor specific CD8^+^ T cells. C57BL/6 mice were implanted with B16-OVA subcutaneously. Two weeks after tumor injection, mice were injected with both IP anti-CD45.2 BB700 antibody and IV anti-CD45.2 AF488 (Fig. 5A). One day after antibody injection, mice were euthanized, and T cell recruitment was assessed by flow cytometry. Using a p15e_604-611_/H-2K^b^ tetramer, a peptide component of a murine endogenous retrovirus protein that is commonly found in mouse cancer cell lines^20^, we found that the IP anti-CD45.2 BB700 identified roughly 7-fold more tumor-specific CD8^+^ T cells that were recently recruited compared to IV anti-CD45.2 AF488 (Fig. 5B). Moreover, when we broadened our analysis to examine all PD-1^+^ CD8^+^ T cells, which likely encompasses the majority of tumor reactive CD8^+^ T cells, we found a similar increase with IP anti-CD45.2 BB700 (Fig. 5C). Finally, analysis of PD-1^-^ CD8^+^ T cells also revealed that IP anti-CD45.2 BB700 labeled significantly more CD8^+^ T cells than IV anti-CD45.2 AF488 (Fig. 5D). These data showing PD-1^-^ CD8^+^ T cells migrate to tumors at a faster rate than PD-1^+^ CD8^+^ T cells are consistent with our prior work in an autochthonous lung cancer model^7^ as well as previous work using photoconvertible alleles (e.g. Kaede) to look at lymphocyte turn over in other transplantable tumors^21,22^. Collectively, these data provide proof of concept that IP anti-CD45.2 BB700 better identifies recently recruited immune cells to tissue, likely due to the increased active labeling time in vivo.

**Fig 5.**
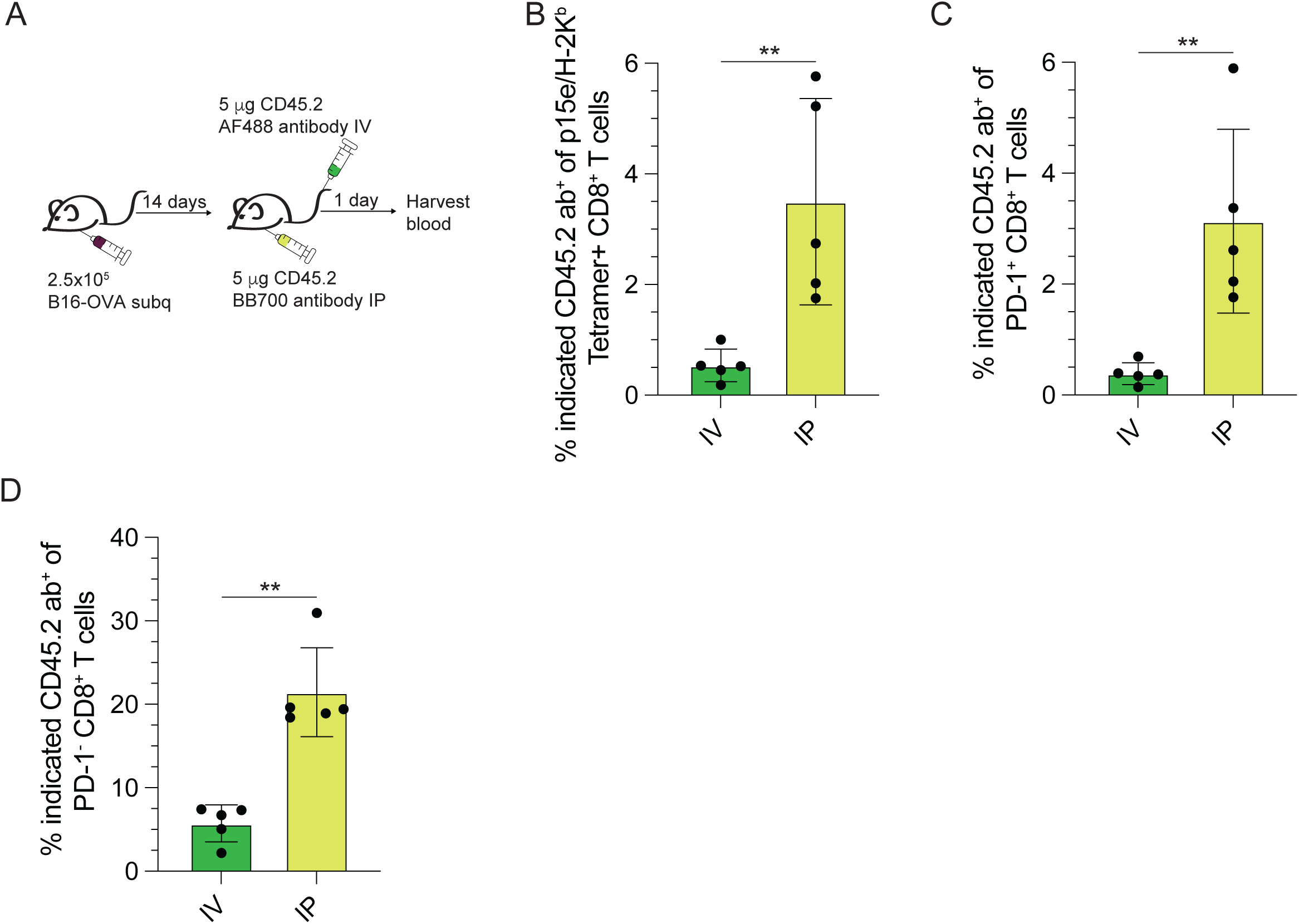
IP anti-CD45.2 antibody detects a great fraction of recruited CD8^+^ T cells to tumor. C57BL/6 mice were subcutaneously injected with 2.5×10^5^ B16-OVA melanoma cells. Two weeks after tumor implantation, tumor bearing mice were co-injected IV anti-CD45.2 AF488 and IP anti-CD45.2 BB700. One day after antibody injections, mice were euthanized and processed for flow cytometry. (A) Experimental schematic. (B-D) Percent positive IP anti-CD45.2 BB700 or IV anti-CD45.2 AF488 on (B) tumor specific p15e/H-2K^b^ tetramer^+^ CD8^+^ T cells, (C) PD-1^+^ CD8^+^ T cells, and (D) PD-1^-^ CD8^+^ T cells. Each dot represents a mouse. Data shown are representative of 2-3 independent experiments, with a total of 6-8 mice per group. Error bars represent SD. Student’s T test was performed to determine statistical significance. **=p<0.01.

## DISCUSSION

Leukocytes must migrate throughout the host to provide on demand protection from perturbations to homeostasis^1,4^. While leukocyte trafficking is important for the immune response in most contexts, the rate, exact mechanisms, and differential contributions of recruited immune cells remains unknown. Recently our group and others described a technique that uses IV injection of anti-CD45.2 antibodies to label circulating leukocytes in the blood before they arrive in tissues, which allows for granular timestamping of leukocytes trafficking to tissues^5–8^. However, we have found that IV anti-CD45.2 antibody was poorly suited for tracking leukocyte migration for cells that were slowly entering tissue. In our work here, we describe how changing route of antibody injection and using different fluorophores can substantially improve the ability to identify rare migration events. Specifically, combining IP anti-CD45.2 antibody with the fluorophore BB700 extended the *in vivo* labeling window from 45 minutes to over 6 hours. We expect that this combination of the technique will be broadly useful for observing rarer migration events *in vivo*.

Our work here also examined the rate and intensity of staining blood contiguous compartments, including the splenic red pulp, liver, and bone marrow. We demonstrate that the liver stains both most rapidly and robustly of these three different organs. Indeed, in both IP and IV anti-CD45.2 antibody injection, the spleen and bone marrow stain more slowly, and the intensity of the staining is much less compared to the liver. This is consistent with previous reports that have looked at both the liver and spleen in short term, *in vivo* labeling experiments^23^. Likely, this decreased rate and intensity of *in vivo* antibody staining is due to a combination of local antibody and immune cell concentration in these different organs. Additionally, the liver is likely one of the first places antibody passes through after injection into the blood, and has fewer immune cells that are not densely packed together. Moreover, because all three of these organs label differently from one another, it also suggests that leukocytes traveling between these locations may differentially stain with *in vivo* antibody. Because of differences in staining intensity between organs, we recommend using this labeling approach to classify leukocytes as being broadly IV antibody positive or negative, rather than making MFI comparisons to quantify antibody abundance between tissues.

Injecting antibody into the peritoneal cavity resulted in the slow release of antibody into the circulation. This result is important for the field beyond our purposes of labeling circulating immune cells. Indeed, many blocking or depleting antibody strategies use IP antibody injection, as this is easier to administer compared to IV injection-based approaches. However, our data raise the important point that injecting a bolus of antibody IP may allow sustained release over the course of multiple hours, as opposed to IV injection where all antibodies would be immediately available in the vasculature. This is a critical point to consider when devising or developing antibody-based therapeutics in pre-clinical mouse models, and one that warrants due consideration prior to initiating experiments.

The *in vivo* anti-CD45.2 antibody staining method will open new avenues of investigation that can be further explored and exploited for a better understanding of what happens when immune cells arrive in tissues. Our original description examining the rate of entry into a mouse model of lung tumors demonstrated the fluidity by which immune cells were continuously pulled into the tumor microenvironment^7^. However, this technique is poised to reveal the transcriptional and epigenetic programming events that occur during leukocyte adaptation in tissues. Our improvement over the original technique using a combination of IP injection and the BB700 fluorophore will help in instances where the population migrating into tissue is only doing so slowly. Moving forward, this technique will need to be further honed, with ideally additional fluorophores to extend labeling in a similar fashion to BB700. Additionally, based on our prior work, there is some labeling of immune cells in the mediastinal lymph node^7^, an important caveat and consideration when using this technique. Future studies may also require additional methodological developments to push past the 3-day antibody labeling limit we have identified.

Taken together, we have demonstrated how both route and fluorophore selection can have significant ramifications for *in vivo* antibody labeling. This methodological refinement will be important beyond tracking leukocyte migration in tumors will aid in better understanding inflammatory diseases including autoimmunity and infection, as well as in examining trafficking events in homeostasis.

## Acknowledgements

We thank the Schenkel and Pauken labs for critical feedback on this manuscript.

## Author contributions

Conceptualization: JMS, KEP

Methodology: JMS, NBH, CI, NK

Investigation: JMS, NBH, CI, NK, TW, TI, AM

Visualization: JMS, NBH, CI

Funding acquisition: JMS

Project administration: JMS Supervision: JMS

Writing – original draft: JMS, KEP, NBH

Writing – review & editing: JMS, NBH, CI, NK, TW, TI, AM, KEP

