## Supplemental Figures for "*In vivo* antibody labeling route and fluorophore dictate labeling efficiency, sensitivity, and longevity" (from another version of this preprint: version 1, 2024-08-10)

### Figure S1

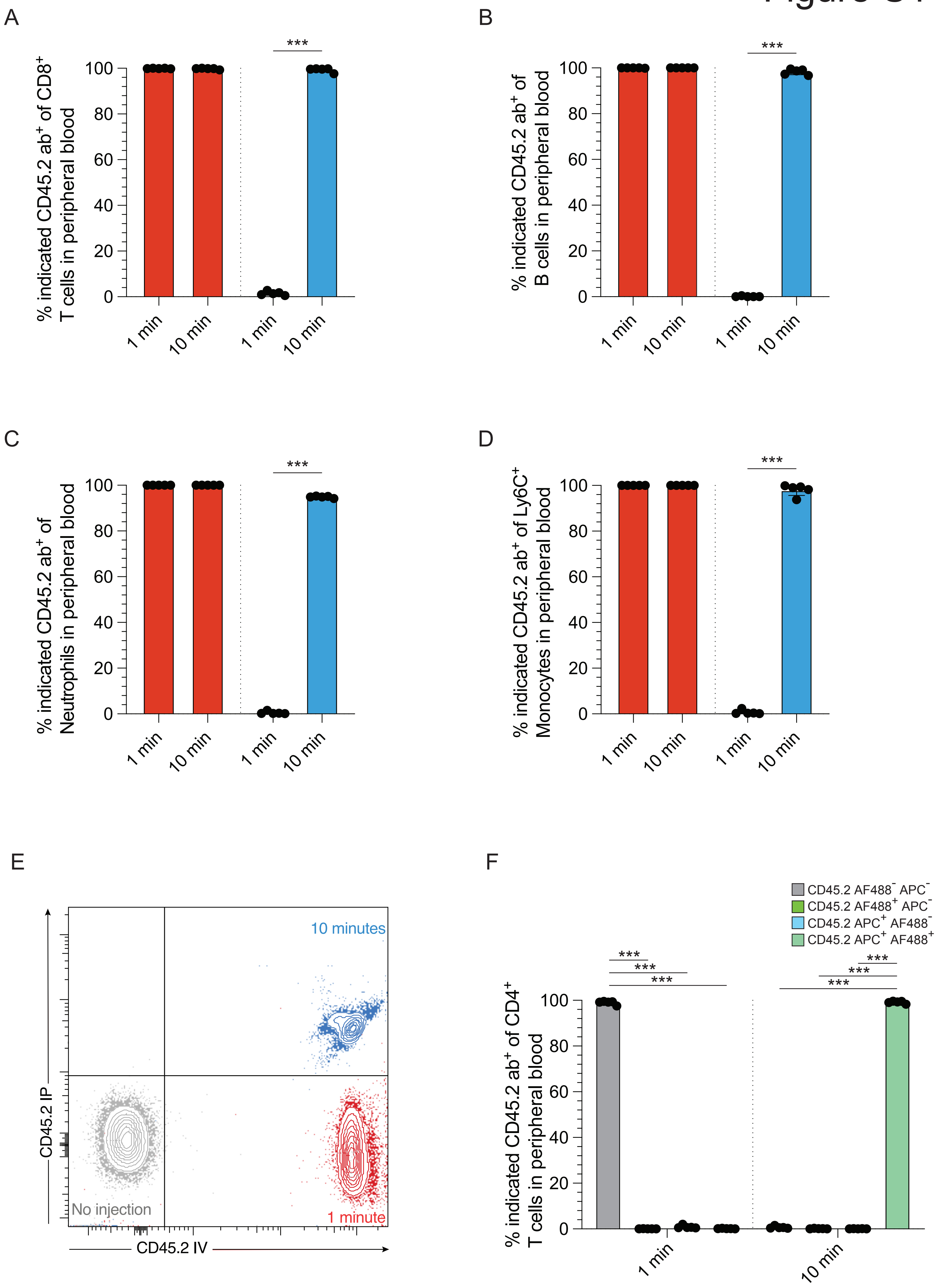
A
% indicated CD45.2 ab+ of CD8+ T cells in peripheral blood
100
80
60
40
20
0
***
1 min
10 min
1 min
10 min
B
% indicated CD45.2 ab+ of B cells in peripheral blood
C
% indicated CD45.2 ab+ of Neutrophils in peripheral blood
D
% indicated CD45.2 ab+ of Ly6C+ Monocytes in peripheral blood
E
10 minutes
No injection
1 minute
CD45.2 IP
CD45.2 IV
F
CD45.2 AF488- APC-
CD45.2 AF488+ APC-
CD45.2 APC+ AF488-
CD45.2 APC+ AF488+
% indicated CD45.2 ab+ of CD4+ T cells in peripheral blood

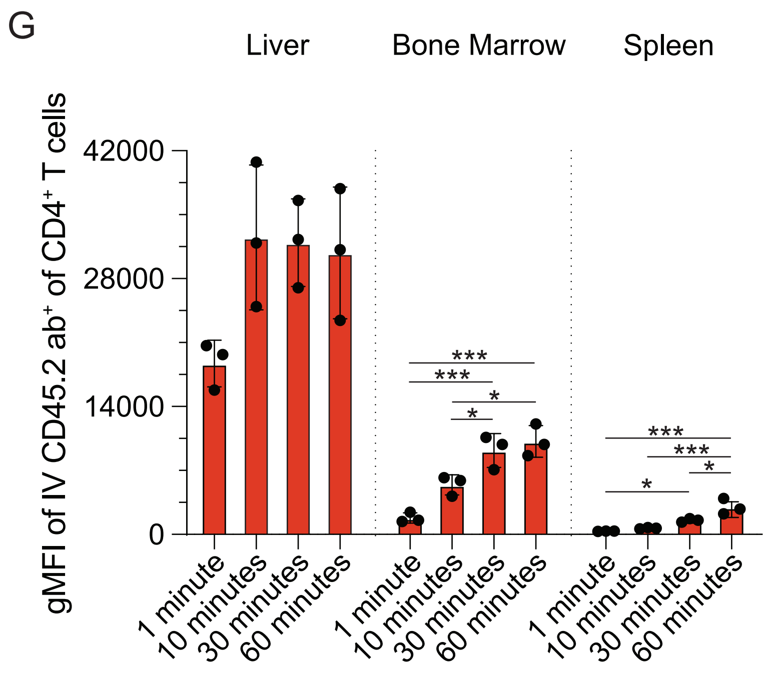
G
Liver
Bone Marrow
Spleen
gMFI of IV CD45.2 ab+ of CD4+ T cells
42000
28000
14000
0
***
***
*
*
***
***
*
*
1 minute
10 minutes
30 minutes
60 minutes

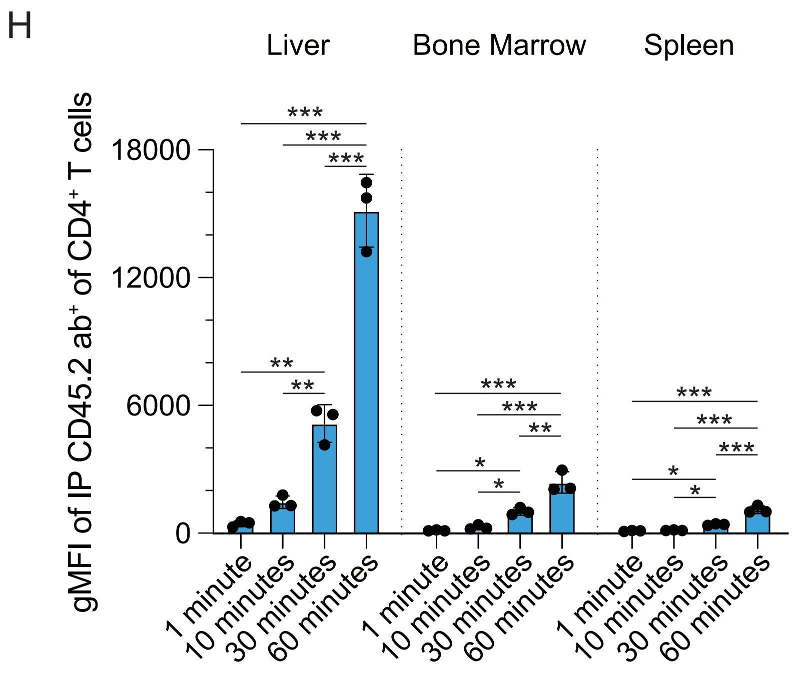
H
Liver
Bone Marrow
Spleen
gMFI of IP CD45.2 ab+ of CD4+ T cells
18000
12000
6000
0
***
***
***
**
**
***
***
**
*
*
***
***
***
*
*
1 minute
10 minutes
30 minutes
60 minutes

**Fig S1. Delayed IP anti-CD45.2 antibody labeling of circulating leukocytes in peripheral and blood contiguous compartments.** (A-E) C57BL/6 mice were co-injected with 5 μg of IV anti-CD45.2 PE and 5 μg of IP anti-CD45.2 APC. Mice were bled one or ten minutes after IP and IV injection and samples were analyzed by flow cytometry. Percentage of (A) $CD8^+$ T cells, (B) B cells, (C) Neutrophils, and (D) Monocytes positive for 1 or 10 minute IV or IP anti-CD45.2 antibody. (E) Representative flow cytometric plot showing staining for IV and IP anti-CD45.2 antibody. No injection control (black), one-minute timepoint (red), and ten-minute timepoint (blue), gated on $CD4^+$ T cells. (F) C57BL/6 mice were co-injected with 5 μg of IP anti-CD45.2 AF488 and 5 μg of IP anti-CD45.2 APC. Mice were subsequently bled either one- or ten-minutes post-injection and analyzed by flow cytometry. (G&H) C57BL/6 mice were co-injected with 5 μg of IV anti-CD45.2 PE and 5 μg of IP anti-CD45.2 APC. Mice were then euthanized either 1, 10, 30, or 60 minutes after injection, and liver, bone marrow, and spleen were harvested and processed for flow cytometry. Geometric mean fluorescence intensity was assessed for (G) IV anti-CD45.2 PE and (H) IP anti-CD45.2 APC on CD4+ T cells in liver, bone marrow and spleen. Each dot represents a mouse. Data shown are representative of 2-3 independent experiments, with a total of 6-8 mice per group. Error bars represent SD. Student's t-test was used for (A-D) and One way ANOVA was performed to determine statistical significance (E-H). $*=p<0.05$,$**=p<0.01$,$***=p<0.001$.

Figure S2

A

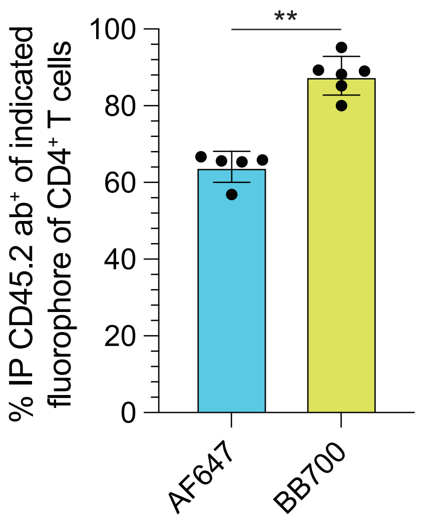

% IP CD45.2 ab⁺ of indicated fluorophore of CD4⁺ T cells
100
80
60
40
20
0
**
AF647
BB700


B

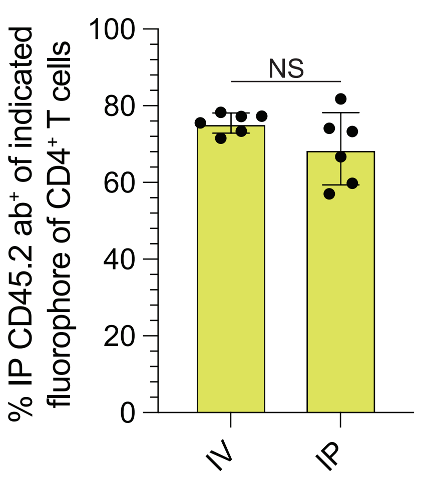

% IP CD45.2 ab⁺ of indicated fluorophore of CD4⁺ T cells
100
80
60
40
20
0
NS
IV
IP

**Fig S2. IP injected anti-CD45.2 BB700 labels more cells than AF647 and is similar in labeling capabilities as IV anti-CD45.2 BB700.** (A) C57BL/6 mice were injected IP with either 5 μg of anti-CD45.2 AF647 or BB700. One day after antibody injection, mice were euthanized and evaluated by flow cytometry. Plotted is the percentage of $CD4^+$ T cells that were positive for IP anti-CD45.2 AF647 or BB700 was quantified. (B) C57BL/6 mice were IV or IP injected with 5 μg of anti-CD45.2 BB700. Three days later, mice were euthanized and staining of $CD4^+$ T cells was evaluated by flow cytometry. Plotted on the graph is percent of $CD4^+$ T cells that are positive for IV or IP anti-CD45.2 BB700. Each dot represents a mouse. Data shown are representative of 2-3 independent experiments, with a total of 6-8 mice per group. Error bars represent SD. Student's t-test was used to determine statistical significance. **=$p<0.01$.
